# PERRC: Protease Engineering with Reactant Residence Time Control

**DOI:** 10.1101/2025.03.02.641063

**Authors:** Sage Nelson, Jokent Gaza, Seyednima Ajayebi, Ronald Masse, Raymond Pho, Cianna Scutero, Samantha Martinusen, Lawton Long, Amor Menezes, Alberto Perez, Carl Denard

## Abstract

Proteases with engineered specificity hold great potential for targeted therapeutics, protein circuit construction, and biotechnology applications. However, many proteases exhibit broad substrate specificity, limiting their applications. Engineering protease specificity remains challenging because evolving a protease to recognize a new substrate, without counterselecting against its native substrate, often results in high residual activity on the original substrate. To address this, we developed Protease Engineering with Reactant Residence Time Control (PERRC), a platform that exploits the correlation between endoplasmic reticulum (ER) retention sequence strength and ER residence time. PERRC allows precise control over the stringency of protease evolution by adjusting counterselection to selection substrate ratios. Using PERRC, we evolved an orthogonal tobacco etch virus protease variant, TEVESNp, that selectively cleaves a substrate (ENLYFES) that differs by only one amino acid from its parent sequence (ENLYFQS). TEVESNp exhibits a remarkable 65-fold preference for the evolved substrate, marking the first example of an engineered orthogonal protease driven by such a slight difference in substrate recognition. Furthermore, TEVESNp functions as a competent protease for constructing orthogonal protein circuits in bacteria, and molecular dynamic simulations analysis reveals subtle yet functionally significant active site rearrangements. PERRC is a modular dual-substrate display system that facilitates precise engineering of protease specificity.

## Introduction

Proteases, enzymes responsible for cleaving proteins through the hydrolysis of peptide bonds, make up approximately 2% of the proteome^1^. These versatile enzymes play a crucial role in numerous essential functions across all living organisms, including immune and inflammatory cell migration and activation^2–4^, wound healing^5–7^, apoptosis^8–10^, and ovulation^11–14^. However, the widespread involvement of proteases in nearly all biological pathways means that any dysregulation can contribute to a broad range of pathologies, including cancers^15–18^, neurodegenerative diseases^19–22^, liver diseases^23–25^, inflammation^26–28^, cardiovascular diseases^29–31^, arthritis^32–34^, and osteoporosis^35–37^. The immense catalytic prowess of these proteases has led to a growing interest in establishing systems to engineer proteases with tailored specificities for proteome editing. Proteases with bespoke substrate specificity could be used as therapeutic agents by degrading disease-associated proteins, ushering in a new wave of precision proteome targeting for disease treatment and cellular reprogramming.

Despite their considerable therapeutic potential, many protease-based drugs struggle to progress beyond clinical trials, largely due to their inherent broad substrate specificity^38,39^. Most proteases exhibit promiscuity, targeting multiple substrates with varying degrees of specificity, which can lead to off-target effects and diminished efficacy^41^. Addressing this challenge requires overcoming the natural multi-specificity of proteases to design variants with tailored substrate preferences. However, successfully altering protease substrate specificity remains a formidable challenge. Often, when proteases are evolved to cleave a desired substrate in the absence of a counterselection substrate, they retain high catalytic activity on their native substrate^40,41^. In fact, often, the engineered protease will have even greater catalytic activity on its native substrate, resulting in an even more promiscuous enzyme^41^. Therefore, it is essential to introduce stringent kinetic competition through the manipulation of counterselection substrate (CS) to selection substrate (SS) ratios^42–47^.

The yeast ER sequestration screening (YESS) system has emerged as a powerful platform for high throughput screening and engineering of enzyme activities^45,48,49,49–52^. Yeast surface display (YSD) offers several advantages over other systems due to its ease of genetic modification, compatibility with flow cytometry analysis, and ability to perform post-translational modifications, including disulfide bond formation^53,54^. The YESS system enables precise control of enzyme:substrate stoichiometries and reaction rates through its transcriptional (promoters of different strengths) and post-translational nodes (ER retention signal (ERS) strengths)^45^. In the present iteration of the YESS system, the CS and SS sequences are located on a single polypeptide, making it impossible to independently change the stoichiometric ratios of CS and SS^45^. This limited functionality makes it impossible to engineer protease substrate specificity under high ratios of CS to SS, which are ideal for guiding evolution campaigns.

Here, we present PERRC, a system for Protease Engineering with ER Reactant Residence Control. PERRC is a yeast ER reactor with orthogonal control of substrate residence times and enzyme:substrate stoichiometries. ERSs dictate how long a cassette remains in the ER^52^. By integrating two substrate cassettes— one with a strong ERS and another with no ERS—into the yeast chromosome, we independently control the residence time of each substrate in the ER. To showcase how the PERRC system can be used to evaluate protease specificity, we tested TEVEp—a TEV protease engineered in a 1:1 CS:SS format to accept a novel P1 residue—and found that the protease loses its orthogonality under the higher CS:SS ratio^48^. We then deployed PERRC to evolve a more specific protease with P1 glutamate specificity. This evolution campaign yielded variant TEVESNp, exhibiting a 64.9-fold specificity switch for glutamate over glutamine at P1, compared to 2.2-fold specificity switch for the parent TEVE enzyme. This specificity switch was also observed in *Escherichia coli*, as confirmed by fluorescence-based protease assays, demonstrating the transferability of TEVESNp’s activity across expression systems. Complementary computational methods based on molecular dynamics (MD) simulations provide additional support for this finding and identifies key conformational changes that drive the shift in binding preferences. Given the significant challenge of achieving such a substantial shift in specificity with a single amino acid change in the substrate sequence, this work underscores the potential of our dual substrate display system as a powerful tool for the precise engineering of protease specificity.

## Results and Discussion

### PERRC is an orthogonal dual substrate display platform with four-color quantification

Engineering protease substrate specificity requires the simultaneous interrogation of protease activities on counterselection and selection substrates^43,44,55–58^. In the standard YESS configuration, both CS and SS are placed on one polypeptide (Figure 1A)^48^. This configuration does not allow differentiation between proteases that cleave both the CS and SS and those that cleave only the CS. Additionally, coupling CS and SS on the same polypeptide restricts the ability to manipulate the CS:SS ratio beyond a fixed 1:1, limiting the flexibility required for precise substrate specificity engineering.

**Figure 1.**
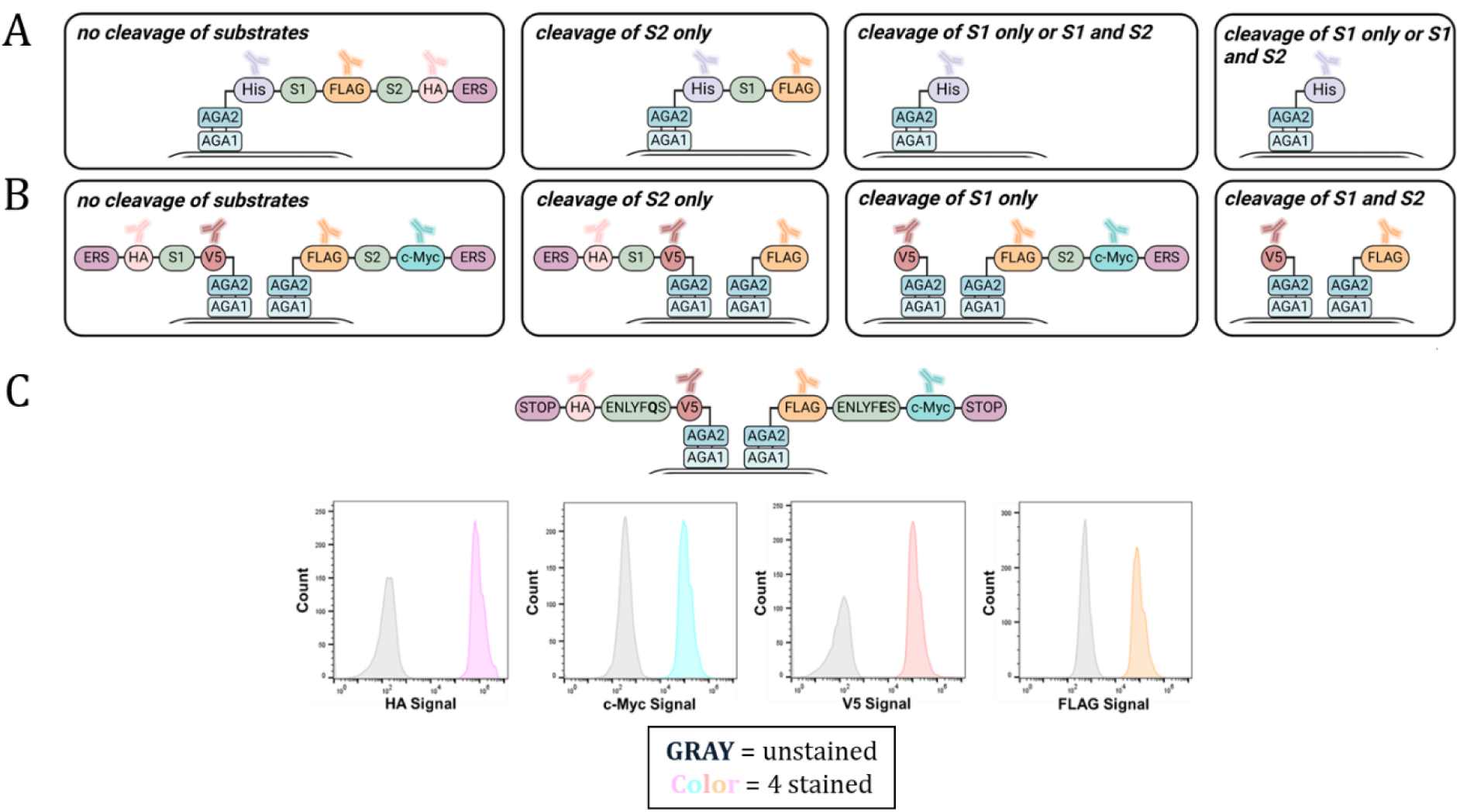
Dual integration enables orthogonal substrate display. (A) In the previous version of YESS CS and SS substrates were within one polypeptide. (B) The new configuration allows us to independently display two or more substrates orthogonally. (C) Flow cytometry histogram plots of the four-stain dual substrate display cassette.

To address these limitations, we developed an orthogonal display system with two AGA2-substrate fusion polypeptides on the yeast surface, enabling independent measurement of cleavage activity for each substrate (Figure 1B). In PERRC, each substrate is flanked with two unique epitope tags, and quantification of substrate processing requires a four-color flow cytometry staining protocol. To achieve this, we integrated two separate AGA2-substrate fusion cassettes at two different loci in the chromosome of EBY100. Namely, an *AGA2-V5tag-Substrate1-HAtag-STOP* was integrated at the *Met15* locus, and an *AGA2-Flagtag-Substrate2-cMyctag-STOP* was integrated at the *HO* locus^59^. Substrate 1 is ENLYFQS (a substrate for WT TEVp), and substrate 2 is ENLYFES (a substrate for the evolved TEV protease TEVEp^48^). We refer to this strain as “Stop-Stop”. Upon induction in galactose medium, surface displayed cassettes were labeled with anti-HA Alexa Fluor® 647, anti-c-Myc Dylight^TM^ 405, anti-FLAG PE, and anti-V5 APC-Cy7 antibodies and visualized via flow cytometry (Figure 1C and Figure S1-S3). Including four distinct fluorescent tags enables the selection of various phenotypes corresponding to the desired proteolytic activity.

PERRC leverages an orthogonal dual substrate display platform with four-color quantification, enabling independent and precise measurement of protease activity on counterselection and selection substrates. By decoupling substrate display and utilizing distinct epitope tags for flow cytometry analysis, PERRC overcomes the limitations of traditional YESS configurations, providing a robust and adaptable framework for engineering protease specificity.

### Enabling independent control of counterselection and selection substrate stoichiometry through endoplasmic reticulum retention sequence manipulations

Appending a strong ERS (WEHDEL) to the C-terminus of an AGA2 substrate fusion slows down surface display, thereby prolonging substrate ER residence time^52^. Therefore, we reasoned that we could manipulate the residence time of two substrate polypeptides independently, for example, by adding a strong ERS to one AGA2 substrate fusion while having no ERS on the other AGA2 substrate fusion. To test this theory, a new dual integration strain was generated^59^. As with the previous “Stop-Stop” strain, we integrated two distinct AGA2-substrate fusion cassettes at separate loci in the EBY100 chromosome. Namely, an *AGA2-V5tag-ENLYFQS-HAtag-WEHDEL* was integrated at the *Met15* locus, and an *AGA2-Flagtag-ENLYFES-cMyctag-STOP* was integrated at the *HO* locus^59^. We refer to this strain as “WEHDEL-Stop” (Figure 2A). We used the previously described “Stop-Stop” strain as a control for quantifying surface display (Figure 2B).

**Figure 2.**
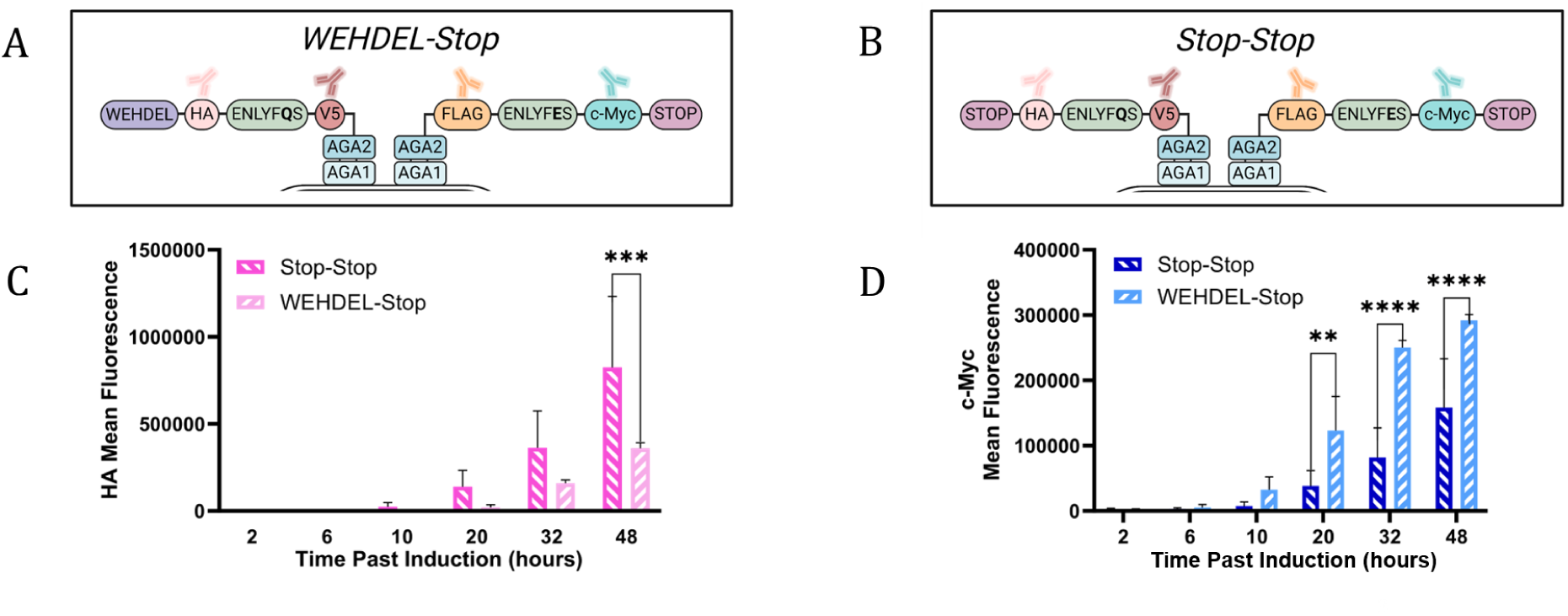
Establishing substrate residence time control in the ER. (A) WEHDEL-Stop cassette where the CS is paired with a strong ERS (WEHDEL), and the SS substrate is paired with no ERS (STOP). (B) Stop-Stop cassette where the CS and SS substrate are both paired with no ERS (STOP). (C) Time course at low temperature (20°C) with dual substrate strains. Flow cytometry bar graph showing the HA mean fluorescence of (A) in light pink and (B) in dark pink. (D) Time course at low temperature (20°C) with dual substrate strains. Flow cytometry bar graph showing the c-Myc mean fluorescence of (A) in light blue and (B) in dark blue. Reactions were run in triplicate. Statistical significance between populations was determined by multiple unpaired t-tests. *p ≤0.05, **p ≤0.01, ***p ≤0.001, ****p ≤0.0001.

Flow cytometry was utilized to assess the influence of ERS on surface display using a low temperature (20°C) 48-hour time course (Figure 2C,D). The low temperature allows for clear observation of gradual shifts in cell surface display. This time course demonstrates that one can control CS to SS ratios through ERS manipulation. As expected, substrates lacking an ERS were transported to the yeast surface faster than those with a strong ERS (WEHDEL) (Figure 2C and Figure S4). At 20 hours post-induction, HA mean fluorescence in the Stop-Stop strain was more than 6-fold higher than in the WEHDEL-Stop strain (Figure 2C). The substrate with the strong ERS, WEHDEL, began to display an HA signal on the surface at 20 hours post-induction. In contrast, the substrate with no attached ERS, Stop, began displaying HA signal on the surface 10 hours post-induction.

Additionally, we observed that AGA1 binding site availability influences surface display. In the Stop-Stop strain, both cassettes leave the ER at a similar rate, leading to increased competition for available AGA1 binding sites on the yeast surface. For example, at the 10 hours past induction time point, there was an over 4-fold greater c-Myc mean fluorescence for the WEHDEL-Stop strain than for the Stop-Stop strain (Figure 2D). This may be because, in the WEHDEL-Stop strain, the WEHDEL cassette stays in the ER longer than the stop cassette. Thus, the Stop cassette may face less competition for AGA1 binding sites. This allows for a higher surface display of the *AGA2-Flagtag-ENLYFES-cMyctag-STOP* cassette, as evidenced by higher c-Myc fluorescence levels in the WEHDEL-Stop strain (Figure 2D). This finding is relevant for protease evolution studies, as a protease that efficiently cleaves ENLYFQS could accelerate WEHDEL ERS removal, potentially increasing competition and reducing surface presentation of the *AGA2-Flagtag-ENLYFES-cMyctag-STOP cassette*.

### Evolving an orthogonal engineered Tobacco etch virus protease with increased specificity for glutamate at P1

Having established orthogonal substrate ER residence time control, we aimed to use it to evolve protease substrate specificity. First, we sought to benchmark the performance of a previously engineered TEV protease, TEVEp^48^. TEVEp is an engineered TEVp that can cleave after a P1 glutamate (ENLYFES). This variant was obtained in the original YESS configuration, where the CS (ENLYFQS) and SS (ENLYFES) were encoded on a single polypeptide with no C-terminal ERS. Furthermore, the TEVp cassette harbored no ERS. Under these conditions, the directed evolution of TEVp yielded a variant that appeared to have switched specificity. However, substrate profiling of TEVEp showed substantial residual activity against the WT sequence^50^. Indeed, in our hands, TEVEp only showed a 2.2-fold preference for ENLYFES over ENLYFQS (Table 1), highlighting the challenge of switching substrate specificities involving a single amino acid change between substrates. By transforming TEVEp into our Stop-Stop strain, we could recapitulate its apparent orthogonality (Figure 3A and C, left pink and blue bars). Under this condition, TEVEp did not cleave large amounts of the ENLYFQS (CS) substrate, resulting in a high HA signal on flow cytometry (Figure 3A and Figure S5). Meanwhile, the c-Myc signal was almost completely lost, indicating high cleavage of the ENLYFES (SS) substrate. However, as expected, when TEVEp was transformed into the WEHDEL-Stop strain, it cleaved both substrates indiscriminately (Figure 3B and C, right pink and blue bars). TEVEp showed an over 6-fold decrease in HA mean fluorescence in the WEHDEL-Stop strain compared to the Stop-Stop strain (Figure 3C). This result further corroborates our hypothesis that a polypeptide with a strong ERS increases its ER concentration relative to one without an ERS.

**Figure 3.**
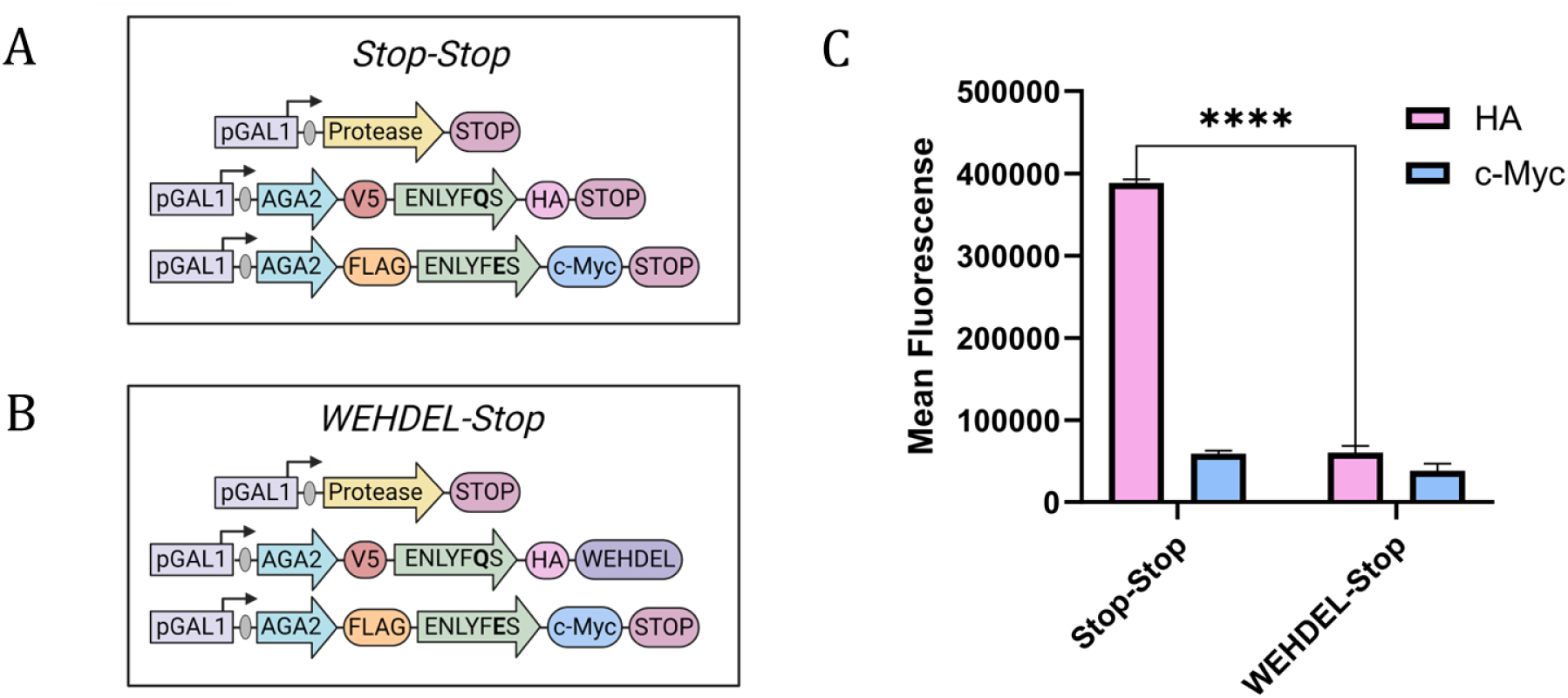
Resident time control in the ER empowers protease engineering under stringent kinetic competition. (A) Stop-Stop cassette where the CS, SS, and TEVEp are all paired with no ERS (STOP). (B) WEHDEL-Stop cassette where the CS is paired with a strong ERS (WEHDEL) and the SS and TEVEp are paired with no ERS (STOP). (C) Flow cytometry bar graph showing HA mean fluorescence (in pink) and c-Myc mean fluorescence (in blue) for both Stop-Stop and WEHDEL-Stop cassettes. Reactions were run in triplicate. Statistical significance between populations was determined by multiple unpaired t-tests. *p ≤0.05, **p ≤0.01, ***p ≤0.001, ****p ≤0.0001.

**Table 1.** Michaelis-Menten Kinetics on TEVEp and TEVESNp on FRET peptide substrates.

| Variant | Mutations | Substrate | $k_{cat}$ (s<sup>-1</sup>) | $K_M$ (mM) | $k_{cat}/K_M$ (mM<sup>-1</sup>·s<sup>-1</sup>) | Fold Increase |
| --- | --- | --- | --- | --- | --- | --- |
| TEVEp | S120R, D148R, T173A, N177K, M218I, S219P | ENLYFES | 173.1 ± 5.6 | 1.82 ± 0.07 | 95.37 ± 4.84 | 2.2 |
|  |  | ENLYFQS | 226.9 ± 24.8 | 5.26 ± 0.84 | 43.17 ± 8.38 |  |
| TEVESNp | **S120C**, D148R, T173A, **N174R**, **N176V**, N177K, **G213A**, M218I, S219P | ENLYFES | 71.41 ± 1.2 | 1.59 ± 0.15 | 44.94 ± 4.40 | 64.9 |
|  |  | ENLYFQS | 6.28 ± 1.1 | 9.08 ± 0.73 | 0.69 ± 0.13 |  |

We hypothesized that TEVEp could be further evolved to exhibit switched specificity if interrogated under a high CS:SS ratio. A variant library of TEVEp with no ERS was generated using error-prone polymerase chain reaction (PCR) (1% error rate) (∼10^8^). This protease library underwent screening against the WEHDEL-Stop dual substrate display system. After inducing protein expression in galactose, 2×10^8^ cells were stained with fluorescently labeled anti-HA-Alexa Fluor® 647, anti-c-Myc FITC, and anti-FLAG PE antibodies. We implemented a two-stage gating strategy. First, we gated on cells with a high HA signal, which corresponds to uncleaved ENLYFQS substrate. From this population, we designed a new gate to collect cells that displayed high PE but low c-Myc signals, which corresponds to cleaved ENLYFES substrate (Figure S6). After three rounds of FACS, we screened 30 single clones and measured their HA and c-Myc signals. We went on to whole plasmid sequence the six best mutants and found that each sequenced mutant had a unique amino acid sequence (Table S1). To further enhance specificity, we performed site saturation mutagenesis at the six identified mutation sites (R105, W143, I166, N174, N176, G213). This DNA library was then transformed into the WEDHEL-Stop strain to make a library (∼10^8^) of TEVEp mutants. As before, the library underwent the high HA/FLAG and low c-Myc gating strategy (Figure S7). After four rounds of sorting, we screened 40 single clones and whole plasmid sequenced the six highest-performing mutants. From the six sequenced clones, five harbored the same mutations: S120C, D148R, T173A, N174R, N176V, N177K, G213A, M218I, and S219P (Table S1). This variant will hereafter be referred to as TEVESNp.

TEVESNp was characterized biochemically using FRET peptides, revealing a striking 64.9-fold preference in catalytic efficiency for ENLYFES over ENLYFQS, compared to 2.2-fold for the parent TEVEp (Table 1 and Figure S8, S9). TEVESNp’s enhanced specificity is largely due to its 36.0-fold lower turnover rate (*k_cat_*) on ENLYFQS compared to TEVEp, significantly reducing off-target cleavage (Table 1). Interestingly, TEVESNp also exhibited a 2.4-fold reduction in *k_cat_* on the ENLYFES substrate relative to TEVEp. This observation aligns with the general principle that evolving a specialist enzyme from a generalist often entails a trade-off in catalytic efficiency to achieve greater specificity. Notably, *K_M_* values remained largely unchanged in TEVESNp, suggesting that the introduced mutations primarily influenced catalytic turnover rather than substrate binding.

### MELDxMD simulations show TEVESNp’s preference for ENLYFESG

Computational tools for studying peptide binding preferences are limited due to the challenges posed by peptide flexibility in predicting binding poses and affinities. For example, docking calculations starting from AlphaFold models (Table S2) yield binding preferences that contradict experiments. To address this, we employ the MELD (Modeling Employing Limited Data)^60,61^ molecular dynamics approach, which has proven effective in predicting relative binding preferences in other peptide–protein systems. MELD leverages multiple copies of the system at varying temperatures and with guiding information, thereby enhancing the sampling of binding/unbinding events and multiple binding modes.

Using a competitive binding framework^62,63^, we investigated the binding preferences of two peptides—ENLYFESG and ENLYFQSG—for both the evolved (TEVESNp) and the parent (TEVEp) enzyme variants (Figure S10). Our simulations reveal distinct differences between the receptors (Figure S11). The TEVEp enzyme accommodates both peptides relatively equally, whereas the TEVESNp receptor has a marked preference for ENLYFESG. Specifically, for TEVESNp, as soon as peptide binding initiates, ENLYFESG dominates across all temperature replicas (Figure 4A). In contrast, for TEVEp, ENLYFESG only outcompetes ENLYFQSG in the lowest temperature replicas, indicating tighter competition between the two peptides.

**Figure 4.**
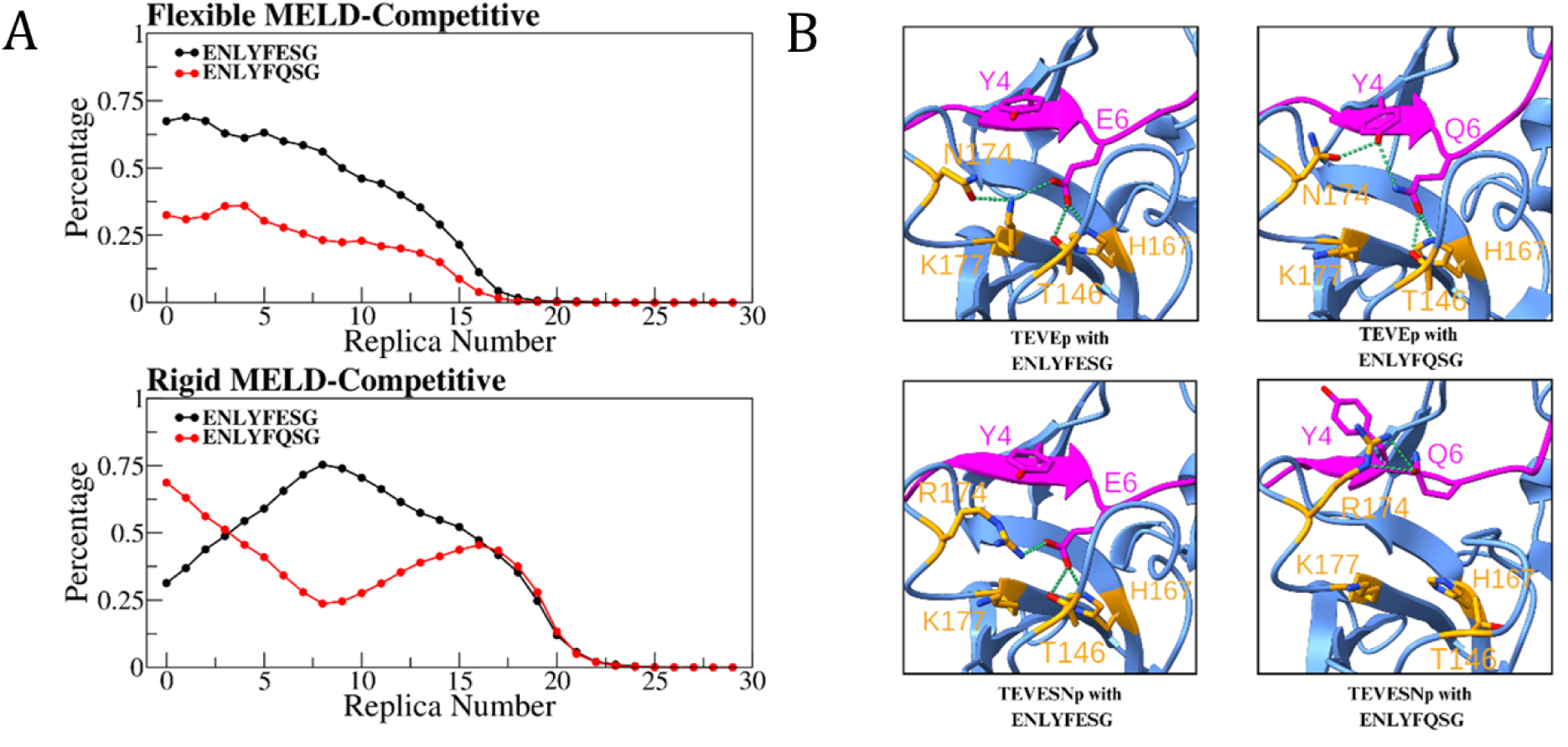
Molecular dynamics simulations of the TEVEp and TEVESNp systems. (A) MELD-competitive simulations of TEVESNp with either a flexible or a rigid protocol for the enzyme. The plots track the population of the bound peptide at each replica, highlighting the clear preference for the ENLYFESG peptide in the TEVESNp enzyme if protein flexibility is taken into account. In the MELD-competitive simulations, multiple binding and unbinding events are expected, with both peptides unbound at the highest replicas. (B) Structural analysis of the centroids of the top clusters from MELD-Bracket simulations. Substrates are shown in magenta, and the enzymes are shown in blue. Important residues in the binding site are represented in stick models.

To probe the role of receptor flexibility, we conducted additional simulations in which the receptor was rigidified while maintaining full peptide flexibility (Figure 4A). Under these conditions, the TEVEp receptor exhibited a reversal in binding preference: ENLYFQSG outcompeted ENLYFESG at all stages. For TEVESNp, rigidification initially led to equal binding of both peptides, with a subsequent enrichment of ENLYFESG at high temperatures and a shift toward ENLYFQSG at lower temperatures.

To clarify these findings, we employed MELD-Bracket—a protocol that improves convergence by seeding each replica with one peptide bound and the other unbound, thereby bypassing the need to sample numerous binding/unbinding events. In MELD-Bracket simulations, we monitor the enrichment of one peptide in the lower replicas and the second one in the higher temperature replicas. These simulations confirm that TEVESNp preferentially binds ENLYFESG (Figure S11), while for TEVEp, the binding preferences remain within error for most replicas, with a slight enrichment of ENLYFQSG only at the lowest temperatures. Taken together, these simulation protocols indicate that TEVEp exhibits tighter competition between the two peptides (i.e., their binding affinities are closer), whereas the evolved TEVESNp displays a clear specificity toward ENLYFESG, in good agreement with experimental observations.

To elucidate the structural basis for TEVESNp’s preference for ENLYFESG, we performed a clustering analysis on the latter halves of the MELD-Bracket trajectories from the five lowest temperature replicas. Visualization of the top cluster centroids (Figure 4B) revealed key differences between the parent and mutant enzymes, as well as between the substrates. In the parent enzyme, both substrates form an extensive hydrogen bond network: E6 and Q6 interact with residues T146 and H167, and residue N174 supports an additional hydrogen bond via its carboxamide side chain—either by interacting with K177 and the negatively charged E6 (in ENLYFESG) or by bonding with Y4’s hydroxyl group, which in turn interacts with Q6 (in ENLYFQSG). When N174 is mutated to R174 in TEVESNp, the longer side chain directly interacts with E6, bypassing the hydrogen bonds with K177 present in the parent enzyme. However, the new hydrogen bond donors in R174 clash with Q6’s amide group, thereby disrupting the interactions with T146 and H167. We surmise that this reconfigured network of interactions reduces the ability of the mutant TEVESNp to interact with ENLYFQSG, leading to an increase of ENLYFESG selectivity.

### TEVESNp’s orthogonality is sustained in a post-translationally controlled protein circuit

To further validate the orthogonality of TEVESNp, we adapted a post-translationally controlled protein circuit in *E. coli* based on cryptic degron-mediated protein degradation^64^. In this system, a fluorescent reporter, green fluorescent protein (GFP), is fused to an engineered degron, which remains inactive until a cognate protease cleaves the upstream recognition sequence, exposing the degron and triggering proteolysis. This approach enables the assessment of protease specificity by linking decrease in GFP fluorescence intensity to protease substrate cleavage efficiency.

To compare the specificity of TEVESNp with WT TEVp and the previously engineered TEVEp variant, we designed two separate reporter constructs in which GFP was preceded by a recognition sequence—either the TEVWTp sequence (ENLYFQ) or the TEVEp/TEVESNp sequence (ENLYFE)— immediately followed by the Y-degron (YLFVQ) (Figure 5A)^64^. Each reporter plasmid was co-transformed into *E. coli* along with a plasmid encoding WT TEVp, TEVEp, or TEVESNp under IPTG-inducible expression. GFP fluorescence was then measured using fluorescence-based degradation assays.

**Figure 5.**
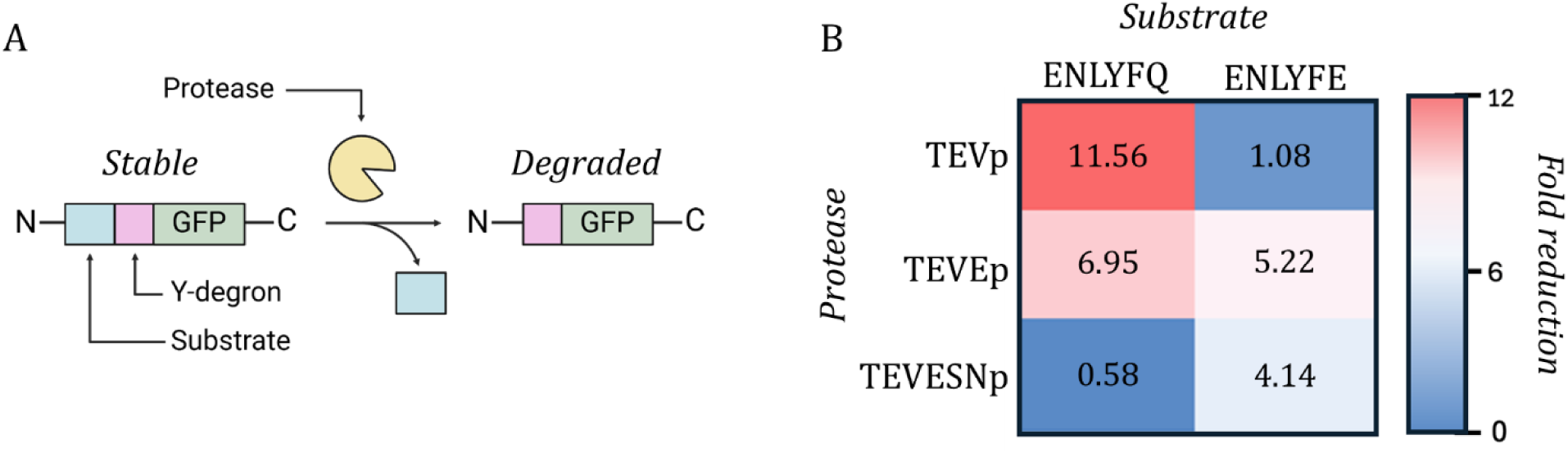
TEVESNp exhibits strong orthogonality in a post-translationally controlled protein circuit. (A) Schematic of GFP degradation through protease cleavage. The blue rectangle is a substrate sequence (ENLYFQ or ENLYFE). The pink square is a N-terminal Y-degron (YLFVQ). The yellow Pac-Mac is the protease (TEVp, TEVEp, or TEVESNp). (B) Orthogonality matrix showing fluorescence fold reduction in cell cultures co-expressing TEVp, TEVEp, or TEVESNp with GFP reporters containing either the ENLYFQ or ENLYFE recognition sequence.

As expected, WT TEVp exhibited an 11.6-fold reduction in fluorescence for the ENLYFQ reporter but only a 1.08-fold reduction for the ENLYFE reporter, confirming its orthogonality. TEVEp displayed an intermediate phenotype, reducing fluorescence by 6.95-fold for ENLYFQ and 5.22-fold for ENLYFE, indicating a lack of orthogonality (Figure 5B). TEVESNp, in contrast, displayed a 4.13-fold reduction in fluorescence for the ENLYFE reporter but only a 0.58-fold reduction—the lowest by far—for the ENLYFQ reporter (Figure 5B). These results align with *in vitro* FRET-based kinetic characterization, reinforcing TEVESNp’s strong specificity for the ENLYFE sequence with minimal cleavage of ENLYFQ. Additionally, TEVESNp exhibited a slightly lower fold reduction for the ENLYFE substrate (4.13-fold reduction) compared to TEVEp (5.22-fold reduction), consistent with its lower *k_cat_* observed in the FRET *in vitro* data (Figure 5B).

These findings demonstrate that TEVESNp achieves highly specific substrate recognition, distinguishing it from TEVEp. Importantly, its specificity is preserved in a post-translationally controlled protein circuit, reinforcing its potential utility for orthogonal protease applications in synthetic biology. The ability of TEVESNp to function with minimal crosstalk in an *E. coli* degradation circuit highlights its robustness and suggests its applicability in modular protein circuits requiring precise protease control.

## Conclusion

The advancement of targeted protease therapeutics relies heavily on developing techniques for engineering protease specificity. The high-throughput, dual-substrate YSD platform presented here allows for versatile enzyme-substrate specificity engineering under varying levels of stringency. An analogy would be offering the protease a plate filled with candy (CS) and selecting only those proteases that exclusively consume spinach (SS). This ER residence time system surpasses traditional methods, such as increasing the copy number of counterselection substrates or fine-tuning promoter and ribosome binding site strengths^65^, in part because differential ER retention has been shown to have a more pronounced impact on enzyme activities^66^. Nonetheless, our system can be augmented with a layer of titratable transcription, offering additional control over selection substrate and enzyme expression levels. Titratable enzyme expression can help manage potential protease toxicity and enable more refined control over catalytic turnovers^67,68^.

Using PERRC, we addressed an oft-unanswered question in protease specificity engineering: whether orthogonal proteases can be evolved to distinguish between highly similar substrate sequences. Early attempts at engineering proteases across such subtle substrate changes—such as converting trypsin into chymotrypsin^69^ and MMP16 into MMP17^70^—yielded promiscuous variants. Not only can TEVESNp differentiate two substrates with only a change at P1, but it also maintains catalytic turnovers comparable to its parent on the counterselection substrate. Considering the challenging nature of this problem, researchers have often found it more sensible to engineer proteases to recognize vastly different substrates _instead40,71,72._

MD simulations played a pivotal role in elucidating the behavior of protease-substrate interactions at the atomic level. These simulations provided critical insights into the conformational flexibility and stability of protease-substrate complexes, deepening our understanding of the forces driving protease-substrate recognition. Furthermore, our findings advance the field of protein circuits. The discovery of a more orthogonal TEVEp variant expands the library of signal processors available for building synthetic protein networks, addressing the current gap in engineered proteases that can be reliably used within these circuits^73^.

Overall, PERRC should allow us to hijack a variety of proteases, leveraging their immense physiological impact to improve human health. Enhanced substrate specificity allows for precise peptide cleavage, improving the effectiveness of biotherapeutics and commercial products while driving progress in proteomics and synthetic biology^39,41,43^. Furthermore, PERCC represents a significant advancement in protease engineering, offering a stringent and modular approach to evolving specificity. Precise control over protease activity opens new avenues for designing tailored proteases with applications in therapeutic development, synthetic biology, and protein circuit engineering. Additionally, the yeast-based platform provides a robust environment for engineering eukaryotic proteases, overcoming a key limitation of bacterial-based platforms^47,74–76^. Together, these innovations expand the toolkit for next-generation protease-based technologies, driving advancements in proteomics, biotherapeutics, and beyond.

## Methods and Materials

### Cassette assemblies

The assembly of cassettes is accomplished through a combination of molecular toolkits MoClo-YTK^77^ (Addgene #1000000061), HiFi DNA Assembly, and Golden Gate Assembly. We have built several receiver plasmids that enable rapid Golden Gate cloning, and such plasmids integrate within the molecular toolkit design process^59^. All substrate cassettes — *AGA2-V5tag-ENLYFQS-HAtag-STOP*, *AGA2-V5tag-ENLYFQS-HAtag-WEHDEL*, and *AGA2-Flagtag-ENLYFES-cMyctag-STOP* — were synthesized by Twist Biosciences. The cassette *AGA2-Flagtag-ENLYFES-cMyctag-STOP* was inserted into the HO-NatR integrative plasmid, while *AGA2-V5tag-ENLYFQS-HAtag-STOP* and *AGA2-V5tag-ENLYFQS-HAtag-WEHDEL* were inserted into the MET15-HYGRO integrative plasmid. These substrate cassettes were assembled using a four-part *BsaI* (NEB, cat# R3733S) Golden Gate reaction comprising pYTK030 (Addgene #65137), the respective substrate cassette, pYTK051 (Addgene #65158), and the corresponding integrative plasmid (Table S3). The TEVE protease cassette (containing mutations listed under the variant name TEV-PE10, previously engineered^48^), includes the TEVE protease sequence followed by a STOP ERS and was synthesized by Twist Biosciences. After digesting the pY3^59^ plasmid with *NheI-HF* (NEB, cat# R3131S) and *XhoI* (NEB, cat# R0146S) to linearize it, the protease cassette was inserted using HiFi DNA Assembly (NEB, cat# E2621S), resulting in pY3-TEVEp (Table S3).

### Yeast strain building

The yeast strains have one substrate integrated at *HO* with nourseothricin resistance (NatR) and the other at *Met15* with hygromycin resistance (HygroR). Construction of yeast strains involved the transformation of substrate containing plasmids into a modified *Saccharomyces cerevisiae* strain (EBY100), which was engineered to express a LexA-hER-haB112 transcription factor^59^. For integration into yeast cells, plasmids were linearized using *NotI* (NEB, cat# R3189S) restriction enzyme digestion, followed by transformation using the Lithium Acetate (LiAc) protocol outlined previously^78^. To enhance recovery, transformed cells underwent an additional 2-hour outgrowth in YPD (yeast extract peptone dextrose) at 30°C following a 42-minute heat shock at 42°C. Transformants were selected on appropriate media (YPD hygromycin for *Met15* integration site and YPD nourseothricin for *HO* integration site) and grown at 30°C for 2-4 days. Mature colonies were then selected and grown in selection media until saturation. The pY3-TEVEp plasmid followed a non-integration transformation into the double-substrate integrated yeast strains, the transformation procedure followed the Frozen-EZ Yeast Transformation II Kit (Zymo Research, cat#T2001), with transformed cells being plated on yeast nitrogen base (YNB) casamino acid (CAA) 2% glucose medium and incubated at 30°C for 2-3 days. Colonies were then selected and grown in YNB CAA 2% glucose 2% raffinose media at 30°C for 16-24 hours until reaching saturation. Selection of transformed cells was performed based on the presence of specific markers: tryptophan for protease-substrate vectors containing a tryptophan gene and leucine for substrate-only vectors containing a leucine gene.

### Yeast and bacterial competent cells preparation

Frozen *E. coli* competent cells were prepared following the protocol outlined in the Mix and Go! *E. coli* Transformation Kit (Zymo Research, cat# T3001) and stored at −80°C. Similarly, yeast competent cells were prepared using the Frozen-EZ Yeast Transformation II Kit (Zymo Research, cat# T2001) and stored at −80°C after preparation.

### Fluorescent labeling, sorting, and cytometry data analysis

Cultures were grown in a deep well 96-well plate (ThermoFisher Scientific, cat# 260252), starting at an OD_600_ of 1 in 0.25 mL. Growth continued until the OD_600_ reached between 2 and 4. Then culture was induced at an OD_600_ of 1 in 0.25 mL of 0.5 in 0.25 mL of minimal galactose-rice media (YNB CAA 2% Galactose). After a 16-hour induction at 30°C shaking at 800 rpm, two million cells were washed in PBS 0.5% BSA (0.5% BSA, Goldbio, cat# 375 9048-46-8) in a round bottom 96-well plate (ThermoFisher Scientific, cat# 174929). Cells were then stained with the appropriate fluorescently labeled antibodies (for 2 million cells: 0.5 µL anti-FLAG PE, 0.5 µL of anti-c-Myc-FITC, 0.5 µL anti-V5 APC-Cy7, and 1 µL anti-HA Alexa 647) (Sigma®’s FLAG® Tag, Biolegend, cat# 637309, anti-HA Alexa Fluor® 647, Biolegend, cat# 682404, Dylight^TM^ 405 AffiniPure^TM^ Goat Anti-Chicken, Jackson ImmunoResearch, cat# 103-475-155, Chicken anti C-MYC epitope tag, exalpha, cat# ACMYC, V5 Epitope Tag Antibody [Allophycocyanin/Cy7], Novus Bio, cat# NB600-381APCCY7). Following stainging, cells were washed and resuspended for cytometric assay (NL Cytek 3000, Cytek Biosciences). Flow Cytometry Standard (FSC) files for each sample assayed using flow cytometry were downloaded into FlowJo (V10.8) for analysis. For data processing, initial yeast cell populations were gated using a forward scatter (FSC-A) (x-axis) vs. side scatter (SSC-A) (y-axis) log-scale plot using a square gate at ≥10^3^ FSC-A and ≥10^3^ SSC-A. After selecting the displaying population, mean fluorescent values of anti-HA, anti-c-Myc, anti-V5, and anti-FLAG signals were extracted. These values were then exported from FlowJo and inputted into GraphPad Software. All quantification measurements are analyzed using a multiple t test with p values corresponding to the following scale: *p ≤0.05, **p ≤0.01, ***p ≤0.001, ****p ≤0.0001.

### Library preparation

The TEVEp error-prone library was prepared with a 1% error rate using Taq DNA polymerase (NEB cat# M0273S) on pY3-TEVEp. After error-prone PCR the sample was incubated with *DpnI* (NEB, cat# R0176S) at 37°C for 1 hour and PCR purified (Zymo Research, cat# D4004). We then conducted a nested PCR on the PCR purified DNA using Taq DNA polymerase (NEB, cat# M0273S). After digesting the backbone, pY3**-**NbLibRec, with *AflII* (NEB, cat# R0520S) and *NsiI-HF* (NEB, cat# R3127S) to linearize it, the error-prone protease was cloned in, and yeast transformation was performed as described previously^79^ (Table S3). Saturated library culture was re-inoculated in a sterile 250-mL flask to an OD_600_ of 0.5 in 50 mL selection media (YNB CAA 2% GLU) and grown at 30°C (250 rpm) until OD_600_ between 2-4 was reached. Once the desired OD_600_ was achieved, the culture was re-inoculated in a sterile 250-mL flask to an OD_600_ of 0.5 in 50 mL induction media (YNB CAA 2% GAL). The induced culture was incubated at 30°C (250 rpm) for 12-15 hours. Post-induction OD_600_ was taken, and the volume of culture required to contain approximately 10x Library-TEVp size was calculated and added to 3 mL PBS 0.5% BSA (0.5% BSA, Goldbio, cat# 9048-46-8). Cells were washed and stained with fluorescently tagged antibodies (Sigma®’s FLAG® Tag, Biolegend, cat# 637309, anti-HA Alexa Fluor® 647, Biolegend, cat# 682404, anti-c-Myc Antibody - FITC Conjugated, Immunology Consultants Laboratory, cat# CMYC-45F-Z) at a concentration of 10^7^ cells/uL. Staining was done for 90 minutes, in the dark, at room temperature. Excess antibody was removed with a second PBS 0.5% BSA wash and resuspended for FACS (BDFACS Melody, BD Biosciences). This same protocol was followed for the preparation of all libraries. After FACS sorting, the DNA was extracted from the confirmed yeast single colonies using a Zymoprep kit (Zymo Research, cat# D2004) and transformed into *E. coli* for Sanger sequencing.

### TEVEp and TEVESNp purification

After digesting pRK793, a gift from David Waugh (Addgene #8827), with *SacI-HF* (NEB, cat# R3156S) and *XbaI* (NEB, cat# R0145S) to linearize it, a gene block from integrated DNA technologies was inserted using HiFi DNA Assembly (NEB, cat# E2621S) in order to change the ENLYFQS cut site to an ENLYFES cut site (Table S3). Sequencing was confirmed with whole plasmid sequencing performed by Plasmidsaurus using Oxford Nanopore Technology with custom analysis and annotation. Subsequently, the new pRK793 with the ENLYFES cut site was digested with *BamHI* (NEB, cat# R0136S) and *XbaI* (NEB, cat# R0145S) to linearize it, then the respective protease sequences were inserted using HiFi DNA Assembly (NEB, cat# E2621S). After sequencing was confirmed with whole plasmid sequencing performed by Plasmidsaurus the plasmids were transformed into BL21 DE3 Codon Plus RIL *E. coli* (Agilent #230245). Using the Ni-NTA FPLC purification protocol similar to that outlined previously^80^, a starting culture (1L inoculation volume) was induced to purify the His-tagged TEVEp. Purified enzyme was aliquoted and flash frozen, with size and purity confirmation conducted using SDS-PAGE (Figure S12).

### TEVp FRET-based activity assays

Fluorogenic peptide substrates: Abz-SENLYFQSG-Lys(DNP) and Abz-SENLYFESG-Lys(DNP) were purchased from Biomatik (Biomatik, Custom Peptide Synthesis). The proteolytic reaction was carried out in 50 mM Tris-HCl (pH 8.0), 1 mM EDTA, and 2 mM DTT at 30°C, with 0.425 µM TEVEp mixed with 0-50 µM respective substrate peptide. Fluoresce was measured via a 320 nm excitation wavelength and a 420 nm emission scan. For each experiment, duplicates of 10 50-uL samples were prepared in the first three columns of a matte black flat bottom 96-well plate (Greiner Bio-One, cat# 655076). Fluorescence intensity (320/405 nm) was measured on a BioTek Synergy Neo2 plate reader at 1 read/15 seconds for 30 minutes. Once all assays were completed, the data was exported into GraphPad Prism (10.1.1) for analysis. Slopes were calculated from the linear regions of each time (x-axis) vs. RFU (y-axis) to get the average reaction velocity, which was subsequently plotted against substrate concentration (on the x-axis) (Figure S11 and Figure S13). Average *K_M_* and V_max_ were extrapolated using a nonlinear regression Michaelis-Menten curve fit, with the error based on the standard deviation of each *K_M_* calculated for each set of substrate concentrations.

### Protease circuit plasmid construction

All vectors used for protease circuit assembly were obtained from the Voigt lab. Modifications were made to the pTEV plasmid (GenBank ID #KX353600) to assemble the pTEVE and the pTEVESN plasmids via HiFi assembly using New England Biolabs NEBuilder HiFi DNA assembly master mix (NEB, cat# E2621S) (Table S3). HiFi compatible gene fragments for TEVEp and TEVESNp were ordered from Twist Biosciences. Lastly, site-directed mutagenesis was conducted on the ptevY_GFP plasmid (GenBank ID# KX353603) to change the TEVp cleavage site from ENLYFES to ENLYFQS using NEBuilder HiFi DNA assembly master mix (cat# E2621S) and overlapping primers (Table S3). The final product was treated with *DpnI* (NEB, cat# R0176S) and was then transformed into NEB 5-alpha competent *E. coli* (NEB, cat# C2987H).

### Plate reader assays for protease circuit characterization

*E. coli* NEB 5-alpha strains harboring protease circuit plasmids were cultured in lysogeny broth (LB) medium supplemented with 100 µg/mL ampicillin at 37°C with continuous shaking at 240 rpm. Cultures were grown overnight, and DNA was extracted using the NEB Monarch Nucleic Acid Purification Kit (NEB, cat# T1110). Similarly, strains containing the Substrate-Degron-GFP construct were cultured under identical conditions but with LB supplemented with 10 µg/mL chloramphenicol. Extracted DNA was whole plasmid sequenced by Plasmidsaurus using Oxford Nanopore Technology with custom analysis and annotation for sequence confirmation.

Chemically competent NEB 5-alpha *E. coli* cells (NEB, cat# C2987H) were co-transformed with all combinations of protease and substrate plasmids. After recovery in Super Optimal Broth (SOB medium), transformants were plated on LB agar supplemented with appropriate antibiotics and incubated at 37°C for 24 hours. Dual plasmid retention was verified before further analysis. Following overnight incubation, cultures and controls were diluted 1:100 into fresh LB medium and incubated for an additional 2 hours. Immediately after, samples were loaded into a 96-well plate and analyzed on a BioTek Synergy Neo2 plate reader.

Each experiment included controls for background fluorescence (media-only wells), baseline substrate fluorescence (substrate-only samples), and non-protease-expressing strains to distinguish circuit activity from autofluorescence or non-specific degradation. All readings were performed in technical and biological triplicates. Fluorescence values were normalized to OD_600_ and background-subtracted before further analysis. Fold change reduction was calculated as the ratio of fluorescence in substrate-only controls to fluorescence in protease-expressing samples.

### Computational methods

MELD (Modeling Employing Limited Data) combines molecular simulations with ambiguous and noisy data through Bayesian inference to identify the most relevant biological states^60,61,81^. MELD uses multiple copies of the system at different temperatures and guiding information in a replica exchange protocol^82^. As a physics-based method, MELD can provide relative binding affinities for peptides binding a receptor^62,63^. Here we take two approaches: in MELD-Competitive, we have two peptides competing for binding the protein receptor, through multiple binding/unbinding events. This method samples multiple binding modes but is challenging to converge. For faster convergence we employ MELD-Bracket, which only samples two state – (1) the target bound to binder A with binder B unbound, and (2) the target bound to binder B with binder A unbound – eliminating the need for repeated binding/unbinding events^83^. In each case, we count for each replica index how often we find peptide A or B in the binding site, giving us a proxy for a relative binding affinity.

### Simulation details

Simulations for MELD-Bracket and MELD-Competitive were initialized from AF3^84^ predictions of the TEVEp and TEVESNp complexes (Figure S13). For MELD-Bracket, two starting states were prepared: (1) enzyme-TENLYFESGT with the TENLYFQSGT peptide shifted 30 Å away, and (2) TENLYFESGT peptide shifted 30 Å away from enzyme-TENLYFQSGT (Figure S10). Simulations used the GBNeck2^85^ implicit solvent model, with proteins modeled using the ff14SB^86^ force field for side chains and ff99SB^87^ for the backbone. MELD simulations employed the OpenMM MD engine and an H,T-REMD protocol with 30 replicas. Temperatures ranged from 300 K at the first replica to 500 K at the highest replica, scaled geometrically. Strong CA-CA distance restraints preserved the enzyme’s folded state based on AF3 predictions and were active across all replicas. Protein flexibility was explored in MELD-competitive simulations by sampling 90% or 100% of these restraints. Restraints were also applied to key protein-peptide interactions from AF3 predictions.

For MELD-Competitive, the initial state included the enzyme with peptides placed 30 Å apart on opposite sides of the binding site (Figure S10). Non-linear scaling was used for binding restraints, transitioning from full strength at the lowest replica to zero at the 18th and above. Distance restraints were employed to both preserve the enzyme’s folded structure and prevent co-binding of binders at low replicas. Both MELD-Bracket and MELD-Competitive simulations ran for 1 µs each. Hydrogen mass repartitioning enabled a timestep of 3.5 fs^88^. In addition to the MELD simulations, we performed 500 ns MD simulations of the enzyme–peptide complexes in explicit water, starting from the MELD-Bracket centroids (Figure 4B). To monitor structural fluctuations, per-residue B-factors were calculated for the latter half of each simulation (Figure S14).

