## Supplemental File for "PERRC: Protease Engineering with Reactant Residence Time Control"

### Supplemental Tables

**Supplemental Table 1.** Detailed list of mutations found in selected variants.

**Supplemental Table 2.** Docking scores of TEVESNp with TENLYFESGT and TENLYFQSGT.

**Supplemental Table 3.** Plasmids and strains used in this study.

### Supplemental Figures

**Supplemental Figure 1.** Representative flow cytometry plots for four color protease activity assays.

**Supplemental Figure 2.** Representative flow cytometry plots displaying anti-c-Myc and anti-V5 fluorescent signals for four color protease activity assays.

**Supplemental Figure 3.** Representative flow cytometry plots displaying anti-FLAG and anti-HA fluorescent signals for four color protease activity assays.

**Supplemental Figure 4.** Representative flow cytometry histogram plots from low temperature time course displaying fluorescent signal of a WEHDEL-Stop cassette containing AGA2-V5-ENLYFQS-HA-WEHDEL and AGA2-FLAG-ENLYFES-c-Myc-Stop.

**Supplemental Figure 5.** Representative flow cytometry plots for TEVEp in the Stop-Stop strain (ENLYFQS with no ERS in blue) and in the WEHDEL-Stop strain (ENLYFES with WEHDEL in red).

**Supplemental Figure 6.** Sorting the error-prone library of TEVEp.

**Supplemental Figure 7.** Sorting the site-saturation mutagenesis library of TEVEp.

**Supplemental Figure 8.** Cleavage of peptide substrates by TEVEp under *in vitro* conditions.

**Supplemental Figure 9.** Cleavage of peptide substrates by TEVESNp under *in vitro* conditions.

**Supplemental Figure 10.** Initial states for MELD simulations.

**Supplemental Figure 11.** Summary of MELD results.

**Supplemental Figure 12.** SDS-Page gels of purified TEV variants.

**Supplemental Figure 13.** AF3 predictions of TEVEp and TEVESNp enzymes with ENLYFESG.

**Supplemental Figure 14.** Per-residue B-factor values and Wormplot representation of the B-factor results.

**Supplemental Table 1**. Evolved TEVEp variants obtained after cell sorting of the initial error-prone PCR library (TEVE1-6p) and evolved TEVEp variants obtained after site-saturation mutagenesis (TEVE7p and TEVESNP) with unique mutations bolded.

| Name | Mutations |
|---|---|
| TEVEp | S120R, D148R, T173A, N177K, M218I, S219P |
| TEVE1p | S120R, D148R, **I166V**, T173A, N177K, M218I, S219P |
| TEVE2p | S120R, D148R, T173A, N177K, **G213S**, M218I, S219P |
| TEVE3p | **R105G**, S120R, D148R, T173A, **N176S**, N177K, M218I, S219P |
| TEVE4p | S120R, D148R, T173A, **N174S**, N177K, M218I, S219P |
| TEVE5p | S120R, D148R, T173A, **N176S**, N177K, M218I, S219P |
| TEVE6p | S120R, **W143R**, D148R, T173A, N177K, M218I, S219P |
| TEVE7p | **R105S**, S120R, D148R, T173A, **N174R**, **N176G**, N177K, M218I, S219P |
| TEVESNp | **S120C**, D148R, T173A, **N174R**, **N176V**, N177K, **G213A**, M218I, S219P |

**Supplemental Table 2.** Docking scores of TEVESNp with TENLYFESGT and TENLYFQSGT.

| Protein | **TENLYFQSGT** | **TENLYFESGT** |
|---|---|---|
| | HPEPDOCK Docking | |
| TEVESNp | -270.207 | -221.593 |
| | SMINA Docking (kcal/mol) | |
| | -10.7 | -10.7 |

**Supplemental Table 3**. Selected plasmids and strains used in this study.

| Strain or Plasmid | Key Characteristics | Plasmid Map (if applicable) |
|---|---|---|
| **Plasmids** | | |
| MET-HYGRO[1] | Integration Site: *MET15*<br>Yeast Marker: HygromycinR | |
| HO-NatR | Integration Site: *HO*<br>Yeast Marker: NourseothricinR | https://benchling.com/s/seq-RUPknqOODiq4TqFVE9dd?m=slm-o8r2zgD2q1ROxjkJJfUv |

| | | |
|---|---|---|
| pY3[1] | Yeast Marker: TRP1 | |
| pY3-NbLibRec | Promoter: *pGAL1*<br>Yeast Marker: TRP1 | https://benchling.com/s/seq-gvaZUxhUiRpdwwrLgY9u?m=slm-7SP8OGFAc7nd2cz3W0RG |
| AGA2-Flagtag-ENLYFES-cMyctag-STOP in HO-NatR | | https://benchling.com/s/seq-wq4va2M7u8gGGJxMfYTR?m=slm-rhZAN0zzAjTcqny4QdD1 |
| AGA2-V5tag-ENLYFQS-HAtag-STOP in Met15-HYGRO | | https://benchling.com/s/seq-fp0ksZ9JSmchG9nrqOpU?m=slm-8YWyaRTYyXkXmyspUj4N |
| AGA2-V5tag-ENLYFQS-HAtag-WEHDEL in Met15-HYGRO | | https://benchling.com/s/seq-rjbRlJ7NxtKVfp98kNl4?m=slm-B0sHEIRYqJRJwXlrLkJB |
| pRK793 with ENLYFES cut site | Bacterial Marker: ChloramphenicolR and AmpicillinR | https://benchling.com/s/seq-3WMFLp6ITp1eXGc1cUF8?m=slm-Im515zi6RmCRsg7PoMMB |
| pRK793 with ENLYFES cut site and TEVESNp inserted | | https://benchling.com/s/seq-CI5oSOZ5lzGLCXrsRgAc?m=slm-KeYuLYNojIkXvA1bCgRb |
| pY3-TEVEp | | https://benchling.com/s/seq-46rGeGK9RrYpqZjG11fJ?m=slm-mszos7lGiZzNXZ9IftB5 |
| pRK793 with ENLYFES cut site and TEVEp Inserted | | https://benchling.com/s/seq-OqJdzaX7S9QKJjvC35Hf?m=slm-szQsTXgOrGkQ7AsAHwLk |
| Protein Circuit ENLYFEY GFP | | https://benchling.com/s/seq-SLApsInquRK9PRZR0Ibx?m=slm-pz7nl1r62NIAGTaEJujF |
| Protein Circuit ENLYFQY GFP[2] | | https://benchling.com/s/seq-7bwsNKRxCxbYYAxrxmT9?m=slm-HKs7OvZGe5pIQPtQEDc3 |
| Protein Circuit TEVEp | | https://benchling.com/s/seq-GYtCMdGekqF1tT9sT0Vp?m=slm-6pS1nCaFX3LOk02PGrIr |

| | | |
|---|---|---|
| Protein Circuit TEVESNp | | https://benchling.com/s/seq-lQaVrIoZHNlI1TO7sj0j?m=slm-qfnyxWVBJU3BBep33FYn |
| FPS Protein Circuit TEVp[2] | | https://benchling.com/s/seq-2G3qUsftoQwEx8MUbJ1l?m=slm-5h0BKyqE0fvPATFXVvvl |
| Integrated DNA technologies gene block for pRK793 cut site change | | https://benchling.com/s/seq-WaW0YiWRm4THdpHRpAdd?m=slm-OpCBgNdK4HFRBmZqVHbz |
| Twist Bioscience "AGA2-Flagtag-ENLYFES-cMyctag-STOP" part | | https://benchling.com/s/seq-mWiqoQBqReAPgpU61dxJ?m=slm-gijOwkucTDaIUVqdwpaU |
| Twist Bioscience "AGA2-V5tag-ENLYFQS-HAtag-STOP" part | | https://benchling.com/s/seq-y9f7Xc75wBrLuYBf7fCI?m=slm-ehNipplbDD80dj3XFjtM |
| Twist Bioscience "AGA2-V5tag-ENLYFQS-HAtag-WEHDEL" part | | https://benchling.com/s/seq-uKGirpbIX5GKMFqVjfNg?m=slm-nLAWKI8OSZf4V9naRbuQ |
| **Strains** | | |
| LLE[1] | *S. cerevisiae* MATa AGA1::GAL1-AGA1::URA3 ura3-52 trp1 leu2-delta200::pACT1-LexA-hER-haB112-ADH1t_pTEF1-ble-TEFt his3-delta200 pep4::HIS3 prbd1.6R can1 GAL | |

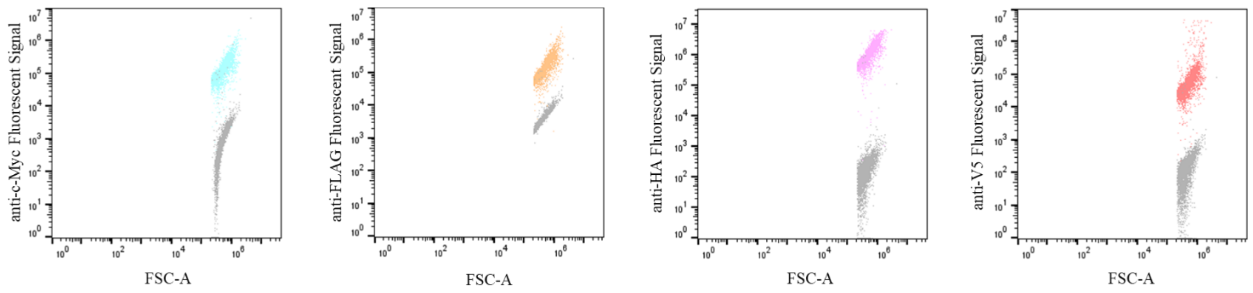


**Supplemental Figure 1.** Flow cytometry dot plots displaying fluorescent signals of four unique fluorescent tags. In this experiments cells stained with four unique fluorescent antibodies were compared to an unstained control. The far-left plot looks at the isolation of anti-c-Myc fluorescent signal (in blue) from four-stained cells and it is compared to unstained cells (in gray). The middle-left plot looks at the isolation of anti-flag fluorescent signal (orange) from four-stained cells and it is compared to unstained cells (in gray). The middle-right plot looks at the isolation of anti-HA fluorescent signal (in pink) from four-stained cells and it is compared to unstained cells (in gray). The far-right plot looks at the isolation of anti-V5 fluorescent signal (in red) from four-stained cells and it is compared to unstained cells (in gray).

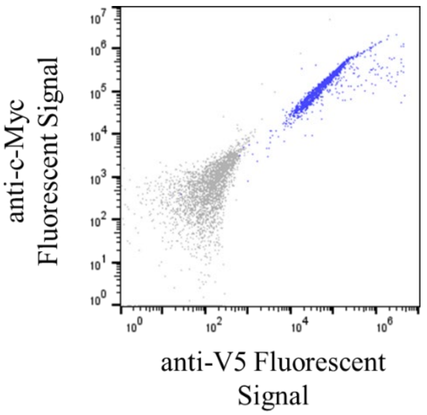


**Supplemental Figure 2.** Flow cytometry dot plot displaying two fluorescent signals from yeast cells stained with four unique fluorescent tags. In this experiments cells stained with four unique fluorescent antibodies were compared to an unstained control. The blue population is c-Myc vs. V5 fluorescent signal from the four-stained cells and it is compared to unstained cells (in gray).

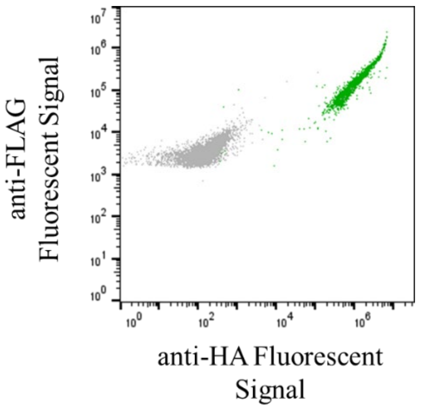


**Supplemental Figure 3.** Flow cytometry dot plot displaying two fluorescent signals from yeast cells stained with four unique fluorescent tags. In this experiments cells stained with four unique fluorescent antibodies were compared to an unstained control. The green population is FLAG vs. HA fluorescent signal from the four-stained cells and it is compared to unstained cells (in gray).

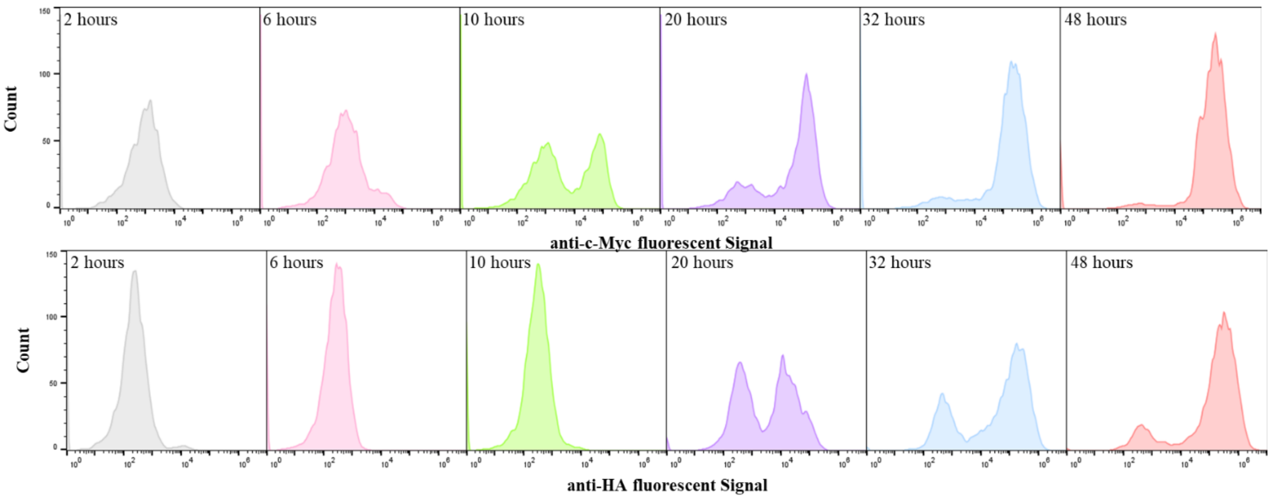


**Supplemental Figure 4.** Flow cytometry histogram plots displaying fluorescent signal of a WEHDEL-Stop cassette containing *AGA2-V5tag-ENLYFQS-HAtag-WEHDEL* and *AGA2-Flagtag-ENLYFES-cMyctag-STOP*. These cells went through a 48-hour time course at a low temperature of 20°C to see the gradual changes in surface display over time. The value in the top left corner of each plot is the number of hours past the initial induction. The top plot is looking at anti-c-Myc fluorescent signal for the WEHDEL-Stop cassette, the c-Myc tag is paired with Stop or no ERS. Surface display occurs with fluorescent signal greater than $10^3$, in the case of c-Myc:Stop this begins to occur at 6-10 hours past induction. The bottom plot is looking at anti-HA fluorescent signal for the WEHDEL-Stop cassette, the HA tag is paired with WEHDEL, a strong ERS. In the case of HA:WEHDEL surface display begins to occur at 20 hours past induction. This assay used technical triplicates (n=3).

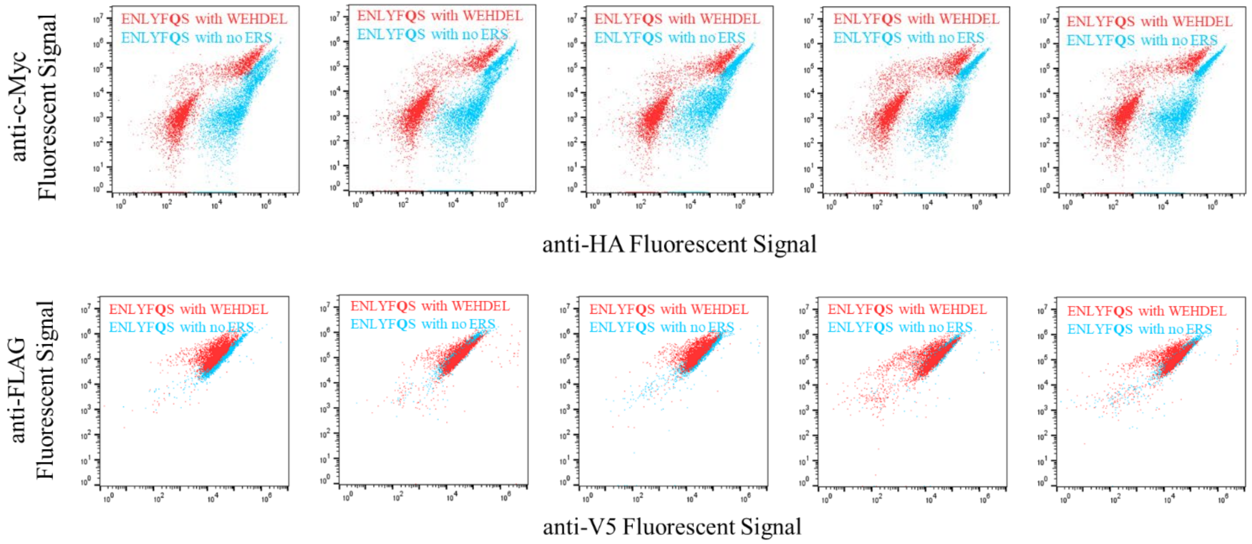


**Supplemental Figure 5.** Representative flow cytometry plots for TEVEp in the Stop-Stop strain (ENLYFQS with no ERS in blue) and in the WEHDEL-Stop strain (ENLYFES with WEHDEL in red). The top flow cytometry plots show c-Myc vs. HA mean fluorescent signal and the bottom plots show FLAG vs. V5 mean fluorescent signal. This assay used technical triplicates (n=3).

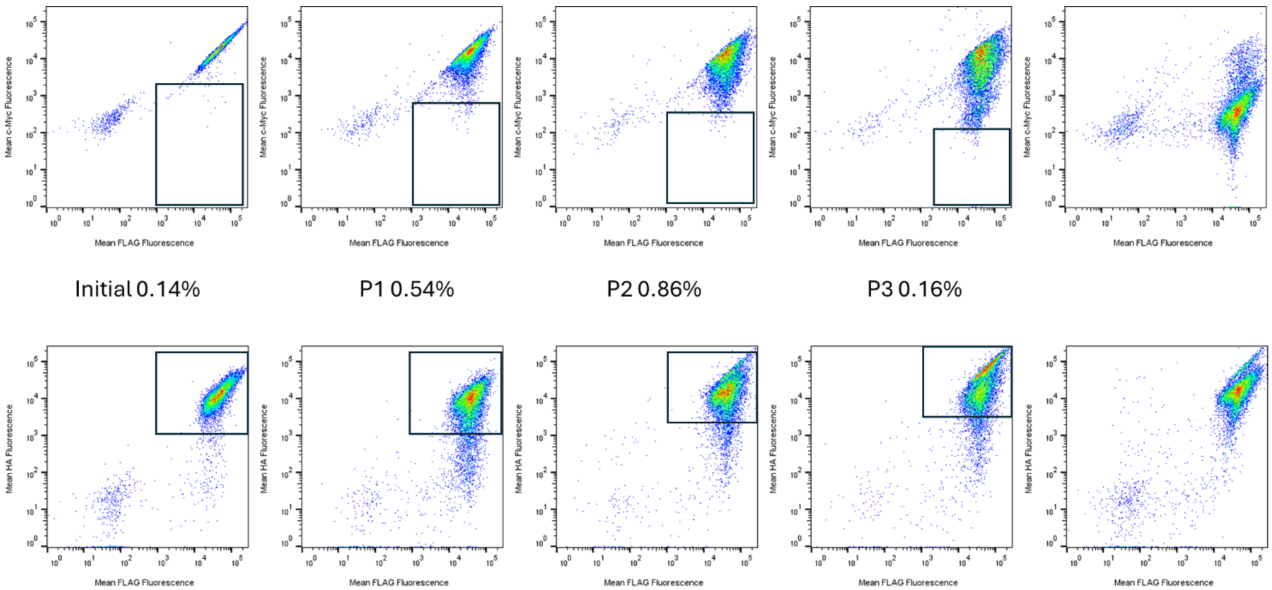


**Supplemental Figure 6.** Sorting the error-prone library of TEVEp. With mean anti-c-Myc fluorescence vs. mean FLAG fluorescence on the top and mean HA fluorescence vs. mean FLAG fluorescence on the bottom.

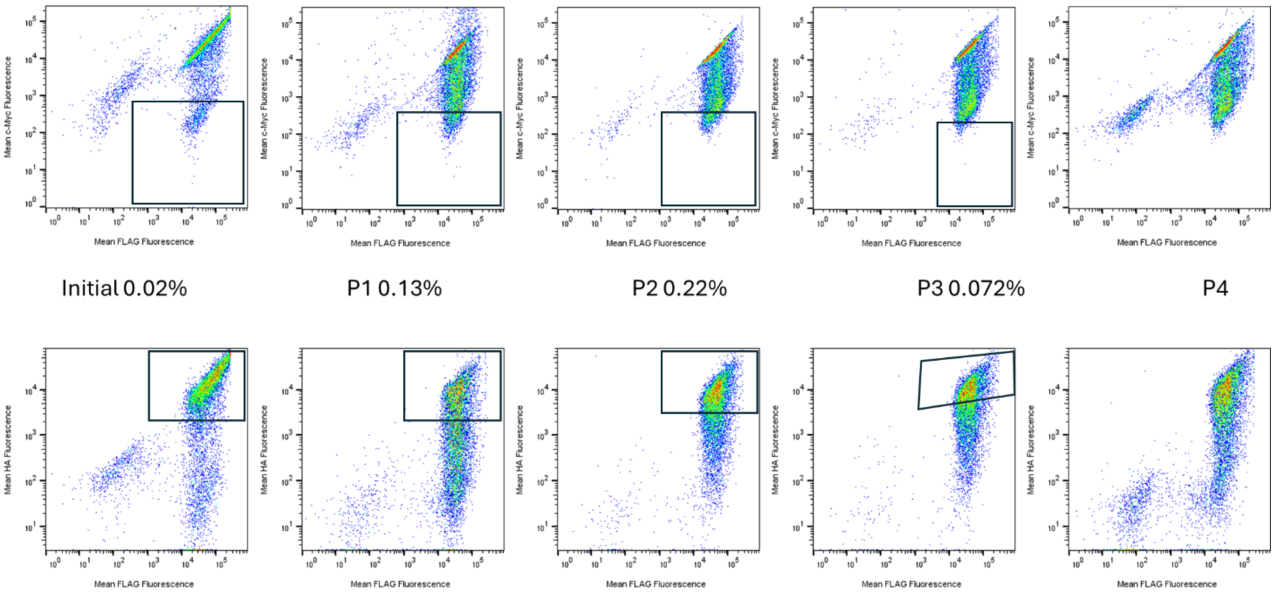


**Supplemental Figure 7.** Sorting the site-saturation mutagenesis library of TEVEp. With mean anti-c-Myc fluorescence vs. mean FLAG fluorescence on the top and mean HA fluorescence vs. mean FLAG fluorescence on the bottom.

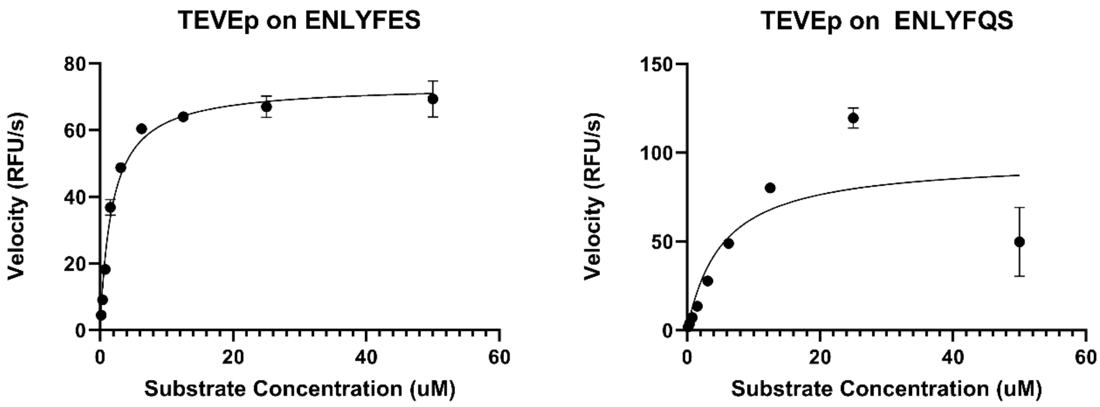


**Supplemental Figure 8.** Cleavage of peptide substrates by TEVEp under *in vitro* conditions. Fluorogenic peptide substrates: Abz-SENLYFQSG-Lys(DNP) and Abz-SENLYFESG-Lys(DNP) were purchased from Biomatik. The proteolytic reaction was carried out in 50 mM Tris-HCl, pH 8.0, 1 mM EDTA, and 2 mM DTT at 30°C, with 0.425 µM TEVEp mixed with 0-50 µM respective substrate peptide. Fluoresce was measure via a 320 nm excitation wavelength and a 420 nm emission scan.

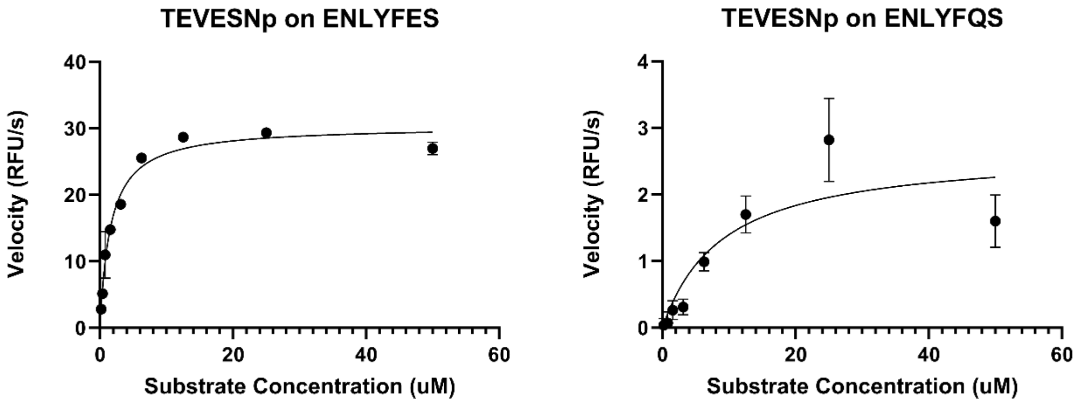


**Supplemental Figure 9.** Cleavage of peptide substrates by TEVESNp under *in vitro* conditions. Fluorogenic peptide substrates: Abz-SENLYFQSG-Lys(DNP) and Abz-SENLYFESG-Lys(DNP) were purchased from Biomatik. The proteolytic reaction was carried out in 50 mM Tris-HCl, pH 8.0, 1 mM EDTA, and 2 mM DTT at 30°C, with 0.425 µM TEVEp mixed with 0-50 µM respective substrate peptide. Fluoresce was measure via a 320 nm excitation wavelength and a 420 nm emission scan.

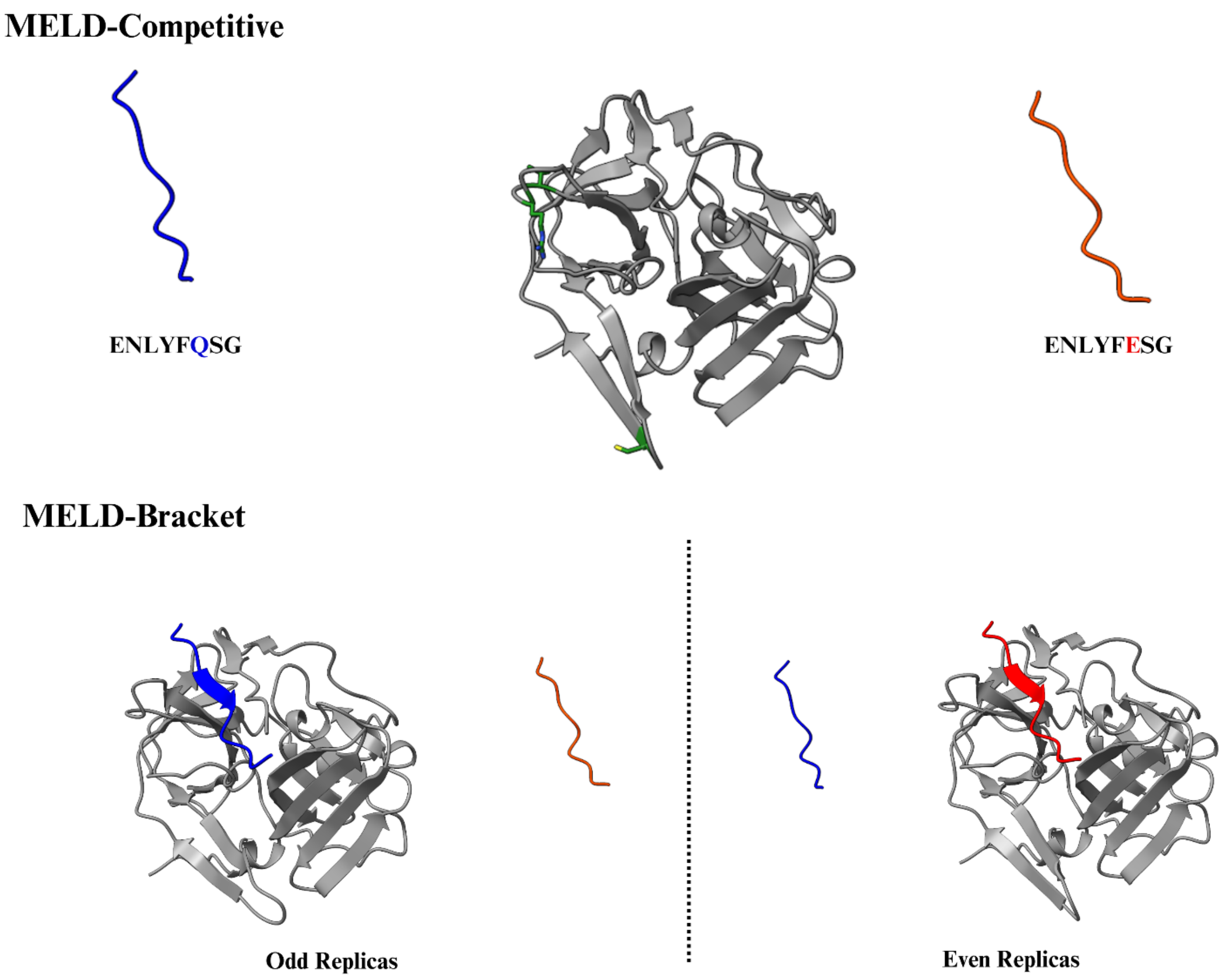


**Supplemental Figure 10.** Initial states for MELD simulations. In MELD-Competitive, both substrates are unbound. Distance restraints are then added to promote binding of only one peptide in low replicas. In MELD-Bracket, two states are periodically distributed along the 30-replica ladder. The relative binding affinities from the two methods are estimated from the population of the bound states at the lowest replicas.

### A. MELD-Competitive (TEVESNp)

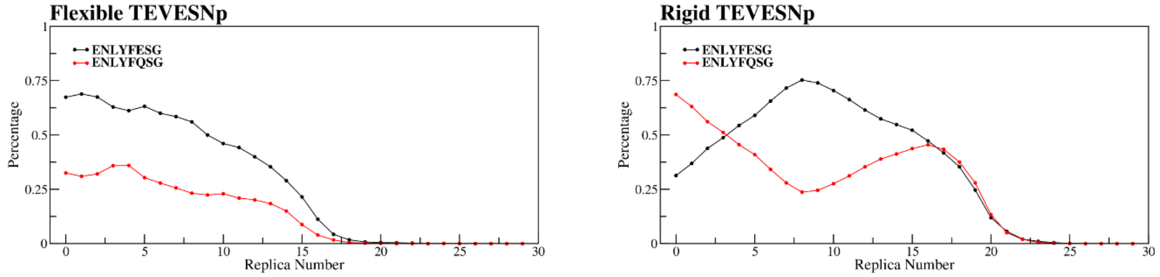


### B. MELD-Competitive (TEVEp)

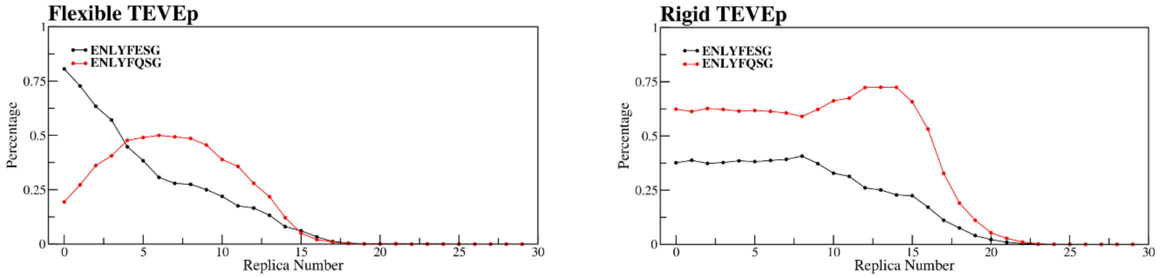


### C. MELD-Bracket

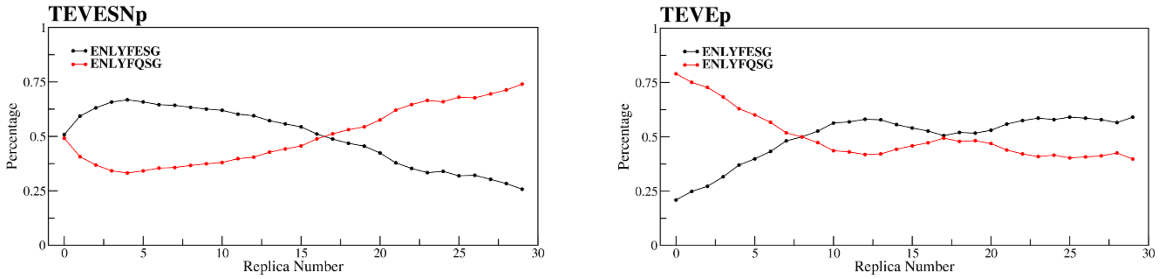


**Supplemental Figure 11.** Summary of MELD results. Showing how much percentage of the simulation (y-axis) each peptide is found at the binding site for each replica index (x-axis).

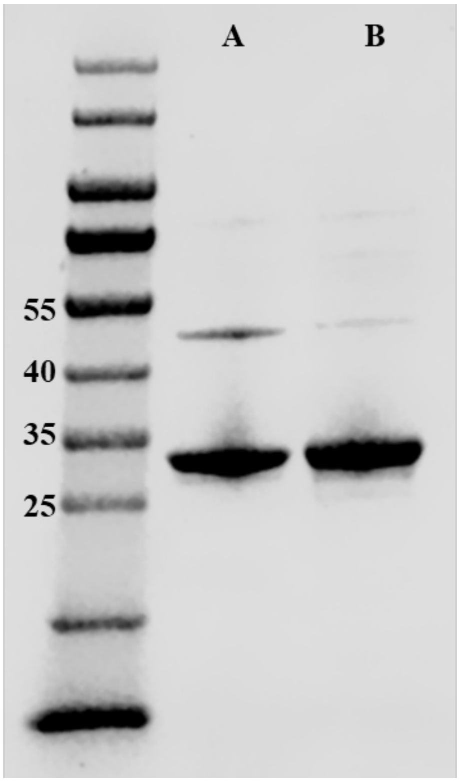


**Supplemental Figure 12.** SDS-Page gels of purified TEVEp (lane A) and purified TEVESNp (lane B).

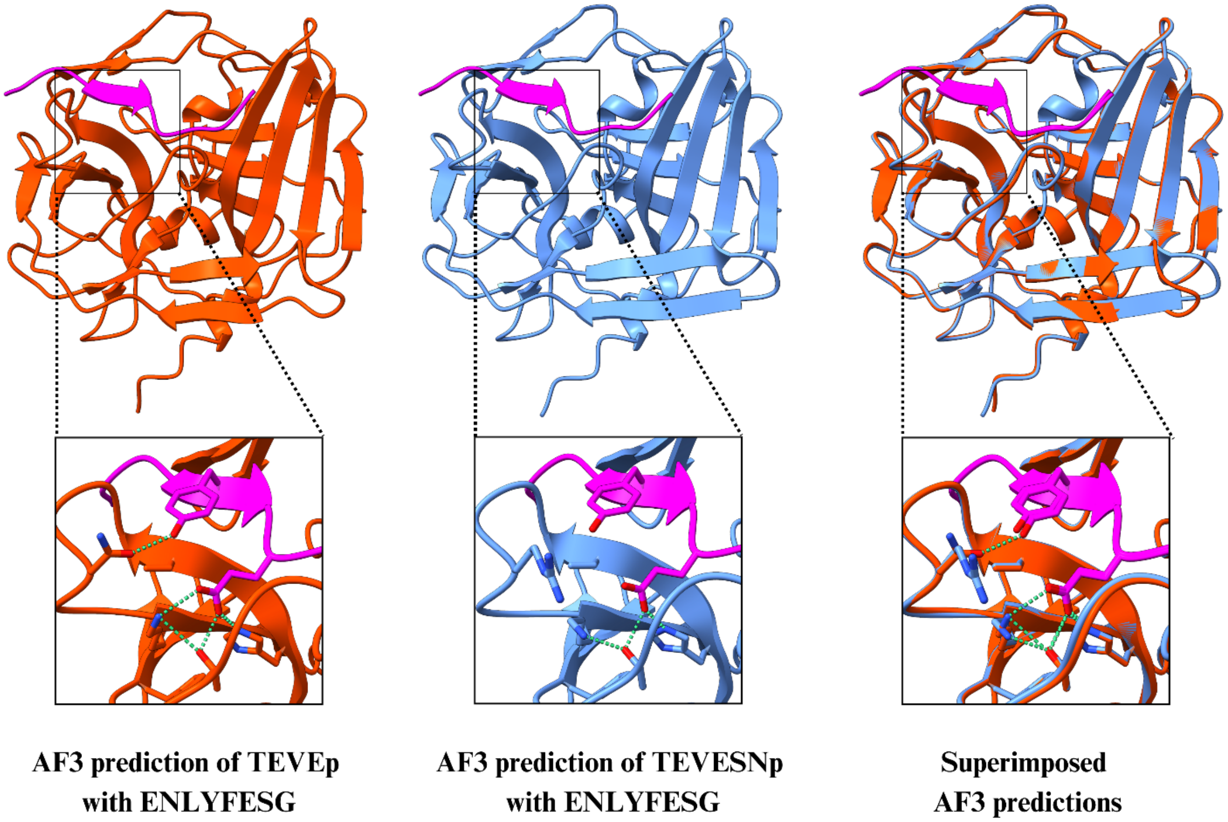


**Supplemental Figure 13.** AF3 predictions of TEVEp and TEVESNp enzymes with ENLYFESG.

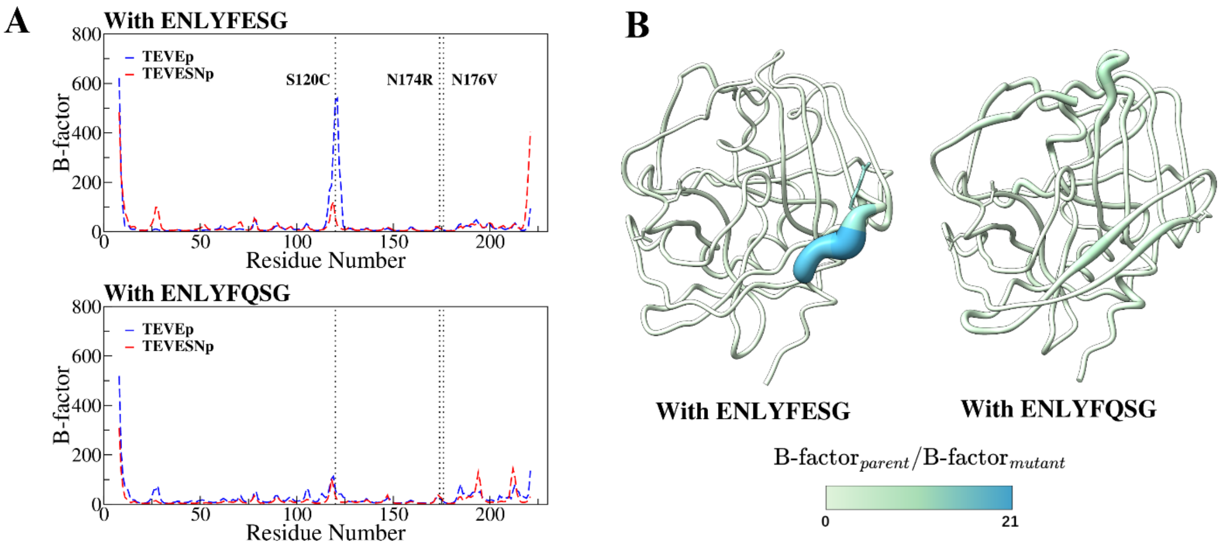

**Supplemental Figure 14.** (A) Per-residue B-factor values from 500 ns explicit MD simulations of the centroids from MELD-Bracket simulations. (B) Wormplot representation of the B-factor results. Residues with increased radii indicate residues with higher fluctuations in the parent enzyme, TEVEp, over the mutant enzyme, TEVESNp. Mutated residues are shown in stick representations. The results show a pronounced spike at residues 119–121 in the TEVEp system when bound to ENLYFESG. This spike disappears when ENLYFQSG is the peptide bound to either the parent or mutant enzyme, and it is reduced in the TEVESNp enzyme. Notably, this region coincides with the distant S120C mutation.

**Supporting Information References**

(1) Martinusen, S. G.; Slaton, E. W.; Nelson, S. E.; Pulgar, M. A.; Besu, J. T.; Simas, C. F.; Denard, C. A. Modular and Integrative Activity Reporters Enhance Biochemical Studies in the Yeast ER. *Protein Engineering, Design and Selection* **2024**, *37*, gzae008. https://doi.org/10.1093/protein/gzae008.

(2) Fernandez-Rodriguez, J.; Voigt, C. A. Post-Translational Control of Genetic Circuits Using *Potyvirus* Proteases. *Nucleic Acids Res* **2016**, *44* (13), 6493–6502. https://doi.org/10.1093/nar/gkw537.
